# Wheat NAC transcription factor *NAC5-1* is a positive regulator of senescence

**DOI:** 10.1101/2024.02.02.578592

**Authors:** Catherine Evans, Sophie Louise Mogg, Charline Soraru, Emma Wallington, Juliet Coates, Philippa Borrill

## Abstract

Wheat (*Triticum aestivum* L.) is an important source of both calories and protein in global diets, but there is a trade-off between grain yield and protein content. The timing of leaf senescence could mediate this trade-off as it is associated with both decline in photosynthesis and nitrogen remobilisation from leaves to grain. NAC transcription factors play key roles in regulating senescence timing. In rice, *OsNAC5* expression is correlated with earlier senescence, but the role of the wheat ortholog in senescence had not been characterised. We verified that *NAC5-1* is the ortholog of *OsNAC5* and that it is expressed in senescing flag leaves in wheat. To characterise *NAC5-1,* we combined missense mutations in *NAC5-A1* and *NAC5-B1* from a TILLING mutant population and overexpressed *NAC5-A1* in wheat. Mutation in *NAC5-1* was associated with delayed onset of flag leaf senescence, while overexpression of *NAC5-A1* was associated with slightly earlier onset of leaf senescence. DAP-seq was performed to locate transcription factor binding sites of *NAC5-1.* Analysis of DAP-seq and comparison with other studies identified putative downstream target genes of *NAC5-1* which could be associated with senescence. This work showed that *NAC5-1* is a positive transcriptional regulator of leaf senescence in wheat. Further research is needed to test the effect of *NAC5-1* on yield and protein content in field trials, to assess the potential to exploit this senescence regulator to develop high-yielding wheat while maintaining grain protein content.

## 1 INTRODUCTION

Wheat (*Triticum aestivum* L.) is the fourth most produced crop globally and production is under pressure from an increasing global population (FAO, 2020). Wheat grain protein is important for both baking quality, as it correlates positively with water absorption and loaf volume (Fradgley et al., 2022), and nutritional value, as wheat contributes 20% of protein in human diets globally (Cauvain, 2012; FAO, 2020). For example, in the UK, criteria for grain to be sold as a ‘Group 1’ breadmaking wheat include a minimum of 13% grain protein content (AHDB, 2023). However, there is a genetic trade-off between yield and protein content, such that past selection for yield has tended to decrease wheat grain protein content (Maeoka et al., 2020; Simmonds, 1995). Novel approaches are needed to understand this trade-off, in order to improve yield while maintaining grain protein content. In addition, development of high protein wheat varieties could ultimately reduce the use of nitrogenous fertiliser currently used to increase wheat grain protein content, which is responsible for nitrous oxide emissions and damage to aquatic environments (FAO, 2020; Yang et al., 2017).

Senescence is the final developmental stage of a wheat plant (Davies & Gan, 2012). The rate of photosynthesis declines during leaf senescence, and delayed leaf senescence correlates with increased yield in cereal crops (Buchanan-Wollaston, 1997; Gregersen, Holm, & Krupinska, 2008; Kichey et al., 2007). However, senescence is required to allow nitrogen remobilisation from leaves to the grain, which accounts for 65-90% of final grain protein content (Bogard et al., 2010; Kichey et al., 2007). Therefore, the timing of leaf senescence is proposed to mediate the trade-off between grain protein content and yield (Thomas & Ougham, 2014). Better understanding of the regulation of leaf senescence may enable the production of higher yielding wheats without a decrease in grain protein content.

The timing of leaf senescence is regulated by a network of transcription factors (Schippers, 2015). In wheat, NAC transcription factor *NAM-1* promotes leaf senescence and grain protein content: introgression of a functional copy of *NAM-B1* led to earlier leaf and peduncle senescence and increased grain protein content in both tetraploid wheat (*Triticum turgidum* ssp. *durum* (Desf.) Husn) and hexaploid wheat *(Triticum aestivum*) (Uauy, Brevis, & Dubcovsky, 2006; Uauy et al., 2006). Delayed leaf senescence and reduced grain protein content were observed in hexaploid RNAi lines with reduced expression of *NAM-A1, NAM-D1* and paralogs *NAM-B2* and *NAM-D2* (Uauy et al., 2006). Similarly, TILLING lines with mutation of either *NAM-A1* in tetraploid wheat (Distelfeld et al., 2012; Pearce et al., 2014) or *NAM-A1* and *NAM-D1* in hexaploid wheat (Avni et al., 2014) and barley near-isogenic lines lacking *HvNAM-1* (Jukanti & Fischer, 2008) displayed delayed senescence and decreased protein content. Mutation of *NAM-A2* and *NAM-B2* in tetraploid wheat delayed flag leaf and peduncle senescence, indicating that *NAM-2*, the paralog of *NAM-1*, also promotes earlier senescence (Borrill et al., 2019). Since the discovery of *NAM-1*, a few other NAC transcription factors have been shown to affect senescence timing in wheat or barley (Christiansen et al., 2016; Harrington, 2019; Zhao et al., 2015), although there are many NAC transcription factors in the wheat and barley genomes that remain to be functionally characterised (Borrill, Harrington, & Uauy, 2017; Murozuka et al., 2018; Vranic et al., 2022).

In rice (*Oryza sativa* L.) NAC transcription factors also act as key regulators of the balance between senescence and grain protein content. The NAC transcription factor *OsNAC5 / NAC071 / ONAC009* (*Os11g0184900*) increases in expression during flag leaf senescence and a high-grain-protein cultivar showed higher expression of *OsNAC5* than a low-grain-protein cultivar, in *O. sativa* ssp. *japonica* and separately in *O. sativa* ssp. *indica* (Sharma et al., 2019; Sperotto et al., 2009). Recombinant inbred lines were developed from *O. sativa* ssp. *indica* cultivars with a range of protein contents; these displayed a correlation between increased grain protein content and increased *OsNAC5* expression in leaves and panicles (Sharma et al., 2019). A T-DNA insertion line with enhanced expression of *OsNAC5* accumulated higher iron and magnesium concentrations in grains, and lower concentrations in leaves (Wairich et al., 2023). These findings suggest that *OsNAC5* may be involved in regulating nitrogen and metal ion remobilization during leaf senescence in rice (Ricachenevsky, Menguer, & Sperotto, 2013). *OsNAC5* has also been associated with ABA-dependent salt, drought and cold stress tolerance (Jeong et al., 2013; Song et al., 2011; Takasaki et al., 2010).

The ortholog of *OsNAC5* in hexaploid wheat, *NAC5-1,* has been reported as the triad *TraesCS4A02G219700*, *TraesCS4B02G098200* and *TraesCS4D02G094400* (Lv et al., 2020). Increased expression of *TraesCS4A02G219700* (*NAC5-A1,* referred to in the cited paper as *TaNAC071-A*), either via transgenic overexpression or a natural promoter insertion allele, was associated with seedling drought tolerance (Mao et al., 2022). This indicates that *TraesCS4A02G219700* shares a role with its ortholog *OsNAC5* in promoting drought tolerance. However, whether *NAC5-A1* shares a role in regulating senescence was not studied. Here we test whether the wheat ortholog of *OsNAC5, NAC5-1,* plays a role in senescence regulation using TILLING mutants, overexpression lines and identification of downstream target genes via DNA affinity purification (DAP-seq).

## 2 MATERIALS AND METHODS

### 2.1 Verification of *OsNAC5* orthologs in wheat

To visualise the orthology of *OsNAC5,* multiple alignments were generated with Clustal Omega 1.2.2 (Madeira et al., 2022). Alignments were visualised using MView, and by generating a consensus tree with the Neighbour-Joining method and 100 bootstrap replicates, in Geneious software (Biomatters, 2022; Madeira et al., 2022). The consensus tree included peptide sequences of *OsNAC5* (Os11t0184900-01 and Os11t0184900-02), the 15 BLASTP hits of *OsNAC5* with lowest E-value from *Triticum aestivum* cv. Chinese Spring (IWGSC RefSeq v1.1) (Appels et al., 2018) and *Oryza sativa* ssp. *japonica* cv. Nipponbare (IRGSP-1.0) (Kawahara et al., 2013; Sakai et al., 2013), and an ortholog of *OsNAC5* in *Physcomitrium patens* (Pp3c13_10800V3_2) as an out-group (Nakashima et al., 2012). The BLASTP search was carried out against the Ensembl Plants database (Yates et al., 2022). Five conserved NAC subdomains assigned to *OsNAC5* were annotated (Kikuchi et al., 2000).

### 2.2 Gene expression data

To assess the pattern of *NAC5-1* expression during flag leaf senescence, transcript levels in wheat flag leaves from 3-26 days after anthesis (DAA) were extracted from RNA-seq data (Borrill et al., 2019). Read counts in transcripts per million (tpm) were extracted via the Wheat Expression Browser and plotted for *NAC5-A1, NAC5-B1* and *NAC5-D1* (Borrill, Ramirez-Gonzalez, & Uauy, 2016).

### 2.3 Selection of TILLING mutant lines

Lines with missense mutations in *NAC5-A1* and *NAC5-B1* were selected from the *Triticum turgidum* ssp. *durum* cv. Kronos TILLING population (Krasileva et al., 2017) (Table 1). Line K2546, with a missense mutation in subdomain iii of the NAC domain of *NAC5-A1,* was crossed with line K2036 and independently with line K3328, with missense mutations in subdomain iv of the NAC domain of *NAC5-B1* (Supplementary Figure 1). Two backcrosses were carried out with non-mutagenized Kronos to reduce the background mutation load. Plants were genotyped at each generation with KASP genotyping as described in the next section. Homozygous double and single mutants were obtained.

**Table 1.** Missense TILLING mutations selected in *NAC5-A1* and *NAC5-B1*. Gene ID from *T. aestivum* RefSeq v1.1 (Appels et al., 2018). Variant ID comprises the wheat line containing the mutation and the chromosome position of the mutation. Alleles of the transcript and amino acid change are shown. Subdomain shows where in five conserved NAC subdomains the mutation sits. PSSM score indicates the likelihood of a missense mutation in this position of the NAC domain across all NAC transcription factors.

| Gene name | Gene ID | Variant ID | Alleles | Amino acid change | Subdomain of NAC domain | PSSM score |
| --- | --- | --- | --- | --- | --- | --- |
| *NAC5-A1* | *TraesCS4A02G219700* | Kronos2546<sup>a</sup> | G/A | G93E | iii | -8 |
| *NAC5-B1* | *TraesCS4B02G098200* | Kronos2036.<br>Chr4B.103056075 | G/A | A111T | iv | -8 |
| *NAC5-B1* | *TraesCS4B02G098200* | Kronos3328.<br>Chr4B.103056044 | G/A | G121D | iv | -4 |

<sup>a</sup>Variant ID not annotated for *NAC5-A1* in RefSeq v1.1 as mutation was initially identified in a previous gene annotation (TGACv1) (Clavijo et al., 2017).

### 2.4 KASP genotyping

For genotyping of mutations in *NAC5-A1* and *NAC5-B1* in Kronos TILLING lines, homoeolog-specific KASP (Kompetitive Allele-Specific PCR, LGC Biosearch Technologies) primers were designed using Polymarker (Ramirez-Gonzalez, Uauy, & Caccamo, 2015). A HEX tag was added to the wild-type allele (WT) primer and a FAM tag to the mutant allele (MUT) primer. Genotyping was carried out at each generation using PACE mix (3CR Bioscience) and KASP markers (Supplementary Table 1) as described in (Ramirez-Gonzalez et al., 2015).

### 2.5 Wheat transformation

The coding sequence for *NAC5-A1* (*TraesCS4A02G219700.1*) was synthesized by GENEWIZ (Azenta). PCR using Q5 High-Fidelity DNA polymerase (NEB), following the manufacturer’s instructions, was used to add a ribosome-binding site immediately upstream of the methionine start codon along with an in-frame 5’ 3xFLAG tag (primers in Supplementary Table 1). The PCR product was cloned into the gateway entry vector pCR8 and checked using Sanger sequencing. The *NAC5-A1* sequence was then recombined into the Gateway-compatible binary vector pSC4-Act-R1R2 in an LR Clonase reaction (ThermoFisher) to create pMSH30, checked by restriction digest and Sanger sequencing then transferred by electrotransformation to *A.tumefaciens* strain EHA105. Plasmids were reisolated from Agrobacterium cultures and verified by restriction digest prior to use in wheat transformation experiments (Bates, Craze, & Wallington, 2017). *NAC5-A1* was expressed *in planta* from the rice Actin promoter (McElroy et al., 1990). Supplementary Figure 2 shows the pMSH130 T-DNA region.

Wheat transformation experiments were set up with spring wheat cultivar Fielder. Immature seeds were harvested 14-20 days after anthesis and immature wheat embryos isolated, then co-cultivated with *A.tumefaciens* for 2 days in the dark (Ishida et al., 2015). Subsequent removal of the embryonic axis and tissue culture with selection agent G418 was performed as previously described (Risacher et al., 2009). Plantlets were hardened off following transfer to Jiffy-7 pellets (LBS horticulture) and genotyped prior to potting in compost.

### 2.6 Copy number assay

Copy number assay was carried out for the marker gene *nptII* against single-copy gene *GAMYB.* Primers and Taqman probes in Supplementary Table 1 were used at a final concentration of 200nM, otherwise as described in (Milner et al., 2018).

### 2.7 Selection of transgenic lines for *NAC5-A1* overexpression

For selection of transgenic lines, T_0_ plants were grown in a controlled environment and a copy number assay was carried out. Five independent transformants with a single construct insertion (“single-copy”) and 25 with more than four construct insertions (“multi-copy”) were obtained (Supplementary Table 2). T_1_ progeny of five single-copy transformants, six multi-copy transformants and two non-transformed controls were grown in the glasshouse.

### 2.8 qPCR

For quantitative PCR (qPCR), 100mg leaf tissue samples from 2 to 3 week-old wheat plants of T_1_ transgenic lines were snap-frozen in liquid N_2_ and ground with a micro-pestle pre-chilled in liquid N_2_. RNA was extracted using TRI-reagent (Invitrogen), RNA was treated with DNAse 1 (ThermoFisher), and cDNA was synthesised with M-MLV reverse transcriptase (Invitrogen), according to manufacturers’ instructions. Primers were designed to amplify *NAC5-A1,* all homoeologs of *NAC5-1* and the transgenic construct (Supplementary Table 1). Using SYBR Green master mix and QuantStudio 5 (Applied Biosystems), qPCR was run with 5 min at 95°C; 45 cycles of 10sec at 95°C, 15sec at 60°C and 20sec at 72°C; and a melt curve from 60°C to 95°C. Fold change in transcript level was calculated using the Pfaffl method incorporating primer efficiencies, with *ACT2* as the reference gene, normalised against the average of single-copy samples (Pfaffl, 2001; Tenea et al., 2011).

For preliminary qPCR, pooled samples of 0.5cm leaf tissue from all plants of one line were collected in the same tube, and qPCR data were analysed by the ΔΔCt method.

### 2.9 Plant phenotyping

Plant growth conditions are summarised in Supplementary Table 3. *NAC5-1* TILLING lines in cv. Kronos at the BC_2_F_3_ generation and *NAC5-A1* transgenic lines in cv. Fielder at the T_2_ generation were phenotyped.

Senescence traits were measured for the primary tiller of each plant. Heading was scored as the date of complete ear emergence from the flag leaf sheath. Flag leaf chlorophyll content was assessed with a SPAD spectrometer, averaging eight points on the leaf surface (SPAD502, Konica Minolta). SPAD readings were made at heading, 6-8 days after heading, then every 7 days until SPAD value decreased below 10 (Supplementary Table 3). Days from heading to a SPAD value of 30 was calculated from the time-course of SPAD values in days after heading using linear interpolation, the same underlying method as previously used to calculate time to a specific leaf senescence score (Chapman et al., 2021). “25% flag leaf senescence” was scored as 25% leaf yellowing and “100% peduncle senescence” as yellowing of the full circumference of the peduncle (Borrill et al., 2019).

Tiller number and height of the primary tiller were measured at maturity. Grain mass, grain number, thousand grain weight, average grain length, grain width and grain area were measured with a Marvin seed analyser (Marvintech). Grain protein content was measured with a Near Infrared Spectrometer (Perten DA7250), normalised for moisture content.

Boxplots were plotted with R package ggplot2 (Wickham, 2016). Lines show median, boxes inter-quartile range and whiskers the most extreme point within 1.5* inter-quartile range from the box. Each time-point of SPAD time-courses, and all traits in *NAC5-A1* overexpression lines, were analysed by a pairwise Wilcoxon test comparing against the control line. Remaining traits in *NAC5-1* TILLING lines were analysed by ANOVA and post-hoc Tukey tests with the formula “Trait ∼ Row + Block + Genotype”. “Block” represents the block of the randomized complete block design, while “Row” represents the row from centre to edge of glasshouse bench, added as a covariate to account for lights positioned in the centre of the bench.

### 2.10 DAP-seq

Coding sequences of NAC transcription factors *NAC5-A1, NAC5-B1, NAM-A1, NAM-B1, NAM-D1, NAM-A2, NAM-B2,* and *NAM-D2* (Gene IDs in Supplementary Table 4) were codon-optimised to reduce GC content using ATGme (Daniel et al., 2015) whilst selecting codons common in wheat to facilitate translation in an *in vitro* wheat germ system (Alexaki et al., 2019; Clarke & Clark, 2008). Constructs were synthesized in a Gateway-compatible entry vector by Twist Bioscience.

Constructs were cloned into the vector pIX-HALO (obtained from *Arabidopsis* Biological Resource Center) in an LR Clonase reaction (ThermoFisher), and checked using Sanger sequencing by GENEWIZ (Azenta). 100mg samples of flag leaf tissue from *T. aestivum* cv. Cadenza at 7 days after heading (DAH) and 14 DAH were collected in liquid N_2_, freeze-dried overnight and homogenized in a Geno/Grinder (Cole-Palmer). Genomic DNA was extracted using a DNA Plant Mini Kit (Qiagen).

DAP-seq was carried out according to (Bartlett et al., 2017), with the following alterations. DNA was sonicated with a BioRuptor Plus (Diagenode) for 21 cycles at 30s on: 30s off, in 200µl aliquots at 20ng/µl, to achieve fragment sizes of 200-400bp (confirmed by Tapestation). DNA was precipitated for 12h at −70°C. Precipitated DNA was pooled, with equal proportions from 7 DAH and 14 DAH. Adapter ligation and library preparation were carried out using NEBNext Ultra II DNA Library Prep Kit for Illumina (NEB), and adapter ligation was checked using qPCR. Proteins were expressed with TNT SP6 High-Yield Wheat Germ Protein Expression System (Promega) and reactions were incubated for 12h at 25°C with 10µg pIX-HALO expression clone. HaloTag beads (Promega) were aliquoted with 2x buffer volume. Three technical replicates were prepared per transcription factor, and two or three replicates passing quality control were sequenced using Illumina 150bp paired end reads. Input DNA controls consisted of 2% DNA library, omitting bead-binding steps, and pIX-HALO controls consisted of DAP-seq carried out using the empty pIX-HALO vector. Primers used to verify cloning and adapter ligation are in Supplementary Table 1.

### 2.11 DAP-seq data analysis

DAP-seq data was analysed following the pipeline in (Klasfeld, Roulé, & Wagner, 2022). Reads were trimmed with Trimmomatic (v0.39), mapped to *T. aestivum* RefSeq v1.1 (Appels et al., 2018) with bowtie2 (v2.4.1), and filtered with samtools (v1.10) view for MAPQ>30 and to remove unmapped reads and secondary alignments. Duplicates were removed with samtools markdup. Two or three technical replicates of the DAP-seq sample preparation were pooled for peak calling, to maximise read depth. Peaks were called with MACS2 (v2.2.7.1) from pooled reads against pIX-HALO controls, to account for the possibility of binding sites of the HaloTag peptide in the wheat genome. A greenscreen mask to remove artifactual peaks, which can arise from amplification, sequencing and mapping biases, was prepared using three input DNA controls with merge_distance=50,000, according to (Klasfeld, Roulé, & Wagner, 2022). The greenscreen contained 228 regions and masked 85 transcripts. Peaks were filtered with q-value < 1 × 10^-10^ or q-value < 0.01, and the greenscreen mask. Peak sets with q-value < 0.01 and greenscreen mask were used for further analyses.

For datasets with >30 peaks, a *de novo* motif search was carried out against peak sequences with RSAT peak-motif (http://rsat.eead.csic.es/plants/peak-motifs_form.cgi), and motifs were compared against the Jaspar database (core nonredundant plants) (http://jaspar2016.genereg.net/). Candidate target genes were identified as the closest high confidence (HC) gene to each peak with bedtools (v2.29.2) closest.

The following sets of candidate target genes were obtained from published data:

1. Candidate target genes of *NAC5-1, NAM-1* and *NAM-2* homoeologs from a GENIE3 gene regulatory network (Ramírez-González et al., 2018). Genes were extracted from the top 1 million connections in the network.
2. Candidate target genes of *OsNAC5* identified by both ChIP-seq, and RNA-seq of *OsNAC5* overexpression lines (Chung et al., 2018). Wheat orthologs of the rice genes were identified with Ensembl Biomart (Kinsella et al., 2011).
3. Candidate target genes of *NAC5-A1* identified by both DAP-seq and RNA-seq of *NAC5-A1* overexpression lines (Mao et al., 2022). Genes were split into upregulated (up) and downregulated (down) genes in *NAC5-A1* overexpression lines compared to wild-type, under well-watered conditions.

The overlap between sets of candidate target genes was assessed using a jaccard test (Chung et al., 2019). The jaccard statistic computes the intersection divided by the union of gene sets. An expected value for the jaccard statistic under the null hypothesis that gene sets are independent was computed based on the set of all high confidence genes in *T. aestivum* RefSeq v1.1 (Appels et al., 2018). Upset plots were generated with UpSetR (Conway, Lex, & Gehlenborg, 2017).

### 2.12 Accession numbers

Raw data for the DAP-seq can be obtained through BioProject ID PRJEB72016 on the European Nucleotide Archive.

Scripts are available at Github https://github.com/Borrill-Lab/NAC5-1SenescenceDAPseq

The following genes were referred to in this study (Supplementary Table 4).

### 2.13 Supplementary Data

Supplementary Figure 1. Crossing scheme for generation of *NAC5-1* TILLING mutant lines.

Supplementary Figure 2. Schematic of construct for *NAC5-A1* overexpression.

Supplementary Figure 3. *TraesCS4A02G219700* and its homoeologs are the orthologs of *OsNAC5* and are expressed in senescence.

Supplementary Figure 4. Additional traits in double mutants of *NAC5-1*.

Supplementary Figure 5. Selection of *NAC5-1* transgenic lines.

Supplementary Figure 6. Further traits in *NAC5-1* T_2_ transgenic lines.

Supplementary Table 1. Primers used in this study.

Supplementary Table 2. Summary of copy number analysis of *NAC5-A1* T_0_ transgenic plants.

Supplementary Table 3. Plant growth conditions.

Supplementary Table 4. Gene IDs used in this study.

Supplementary Dataset 1. Peaks and closest HC genes to peaks from DAP-seq.

Supplementary Dataset 2. Intersection between closest genes to DAP-seq peaks in this study and published gene sets.

Supplementary Dataset 3. Genes which overlap in two or more gene sets across this study and published gene sets.

Supplementary Dataset 4. Functions of candidate *NAC5-1* downstream target genes.

## 3 RESULTS

### 3.1 *NAC5-1* is the ortholog of *OsNAC5* and shows increased expression during flag leaf senescence

The triad of wheat genes *TraesCS4A02G219700*, *TraesCS4B02G098200*, and *TraesCS4D02G094400* has previously been noted as the ortholog of *OsNAC5/ONAC071* (Lv et al., 2020). In this study, these genes will be referred to as *NAC5-A1* (*TraesCS4A02G219700*)*, NAC5-B1* (*TraesCS4B02G098200*) and *NAC5-D1* (*TraesCS4D02G094400*) (Supplementary Table 4), in accordance with guidelines on wheat gene nomenclature based on homology (Boden et al., 2023). *NAC5-A1* shows 97.3% and 98.5% identity with homoeologs *NAC5-B1* and *NAC5-D1* respectively, and 81.9% identity with *OsNAC5* at the peptide level (Supplementary Figure 3a). *NAC5-1* was the only wheat gene triad to cluster with *OsNAC5* based on peptide alignment of related wheat and rice NAC transcription factors, confirming that *NAC5-1* and *OsNAC5* have 1:1 orthology (Supplementary Figure 3b). To assess the suitability of *NAC5-1* as a candidate senescence regulator, the expression of *NAC5-1* in a time-course of flag leaf senescence was extracted from an RNA-seq dataset (Borrill et al., 2019). *NAC5-A1, NAC5-B1* and *NAC5-D1* increase in expression from 3 days after anthesis (DAA) to 26DAA, with *NAC5-D1* also showing a peak in expression at 15DAA (Supplementary Figure 3c).

### 3.2 Missense mutations in *NAC5-1* are associated with a delay in leaf senescence

In rice, *OsNAC5* expression is positively correlated with earlier senescence. Therefore, to test the hypothesis that loss-of-function of *NAC5-1* will delay senescence, lines with mutations in all copies of *NAC5-1* were developed by crossing together lines from the tetraploid wheat cv. Kronos TILLING population (Krasileva et al., 2017). Missense mutations in the highly conserved NAC domain responsible for both DNA binding and dimerization (Ernst et al., 2004; Welner et al., 2012) were identified in both *NAC5-A1* and *NAC5-B1* (Table 1, Figure 1a). Line K2546, with a missense mutation in subdomain iii of the NAC domain of *NAC5-A1,* was crossed with line K2036 and independently with line K3328, which both had missense mutations in subdomain iv of the NAC domain of *NAC5-B1* (Supplementary Figure 1). Two backcrosses were carried out with wild type Kronos to reduce the background mutation load.

**Figure 1.**
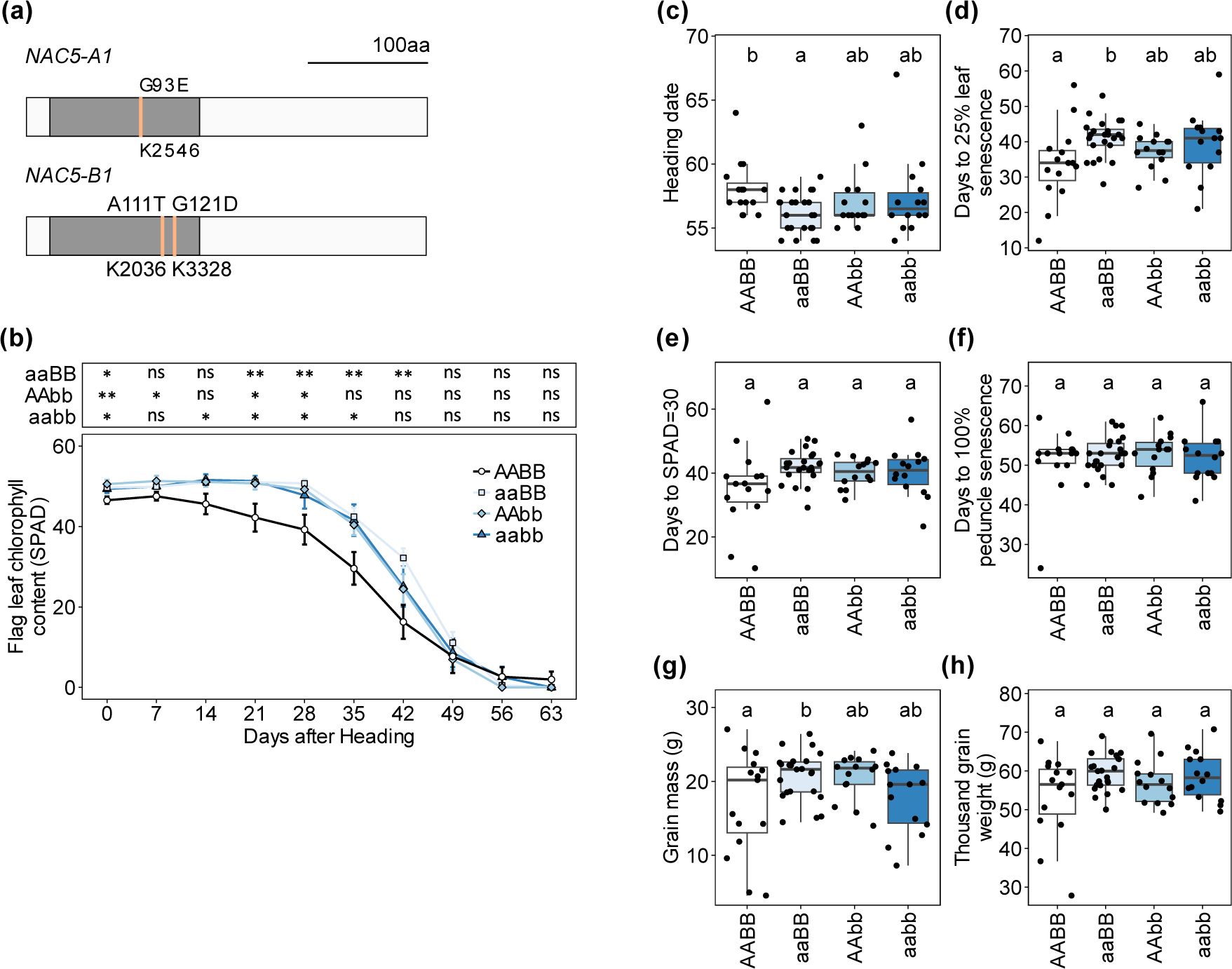
*NAC5-1* TILLING lines retain higher chlorophyll content than controls. (a) Peptide sequences of *NAC5-1*, annotated with selected TILLING mutations in Kronos. Pale lines mark mutations, white boxes show peptide sequence, grey boxes show conserved NAC domain. (b) Flag leaf chlorophyll content (SPAD) in days after heading (DAH) +/− 1 day was compared between wild type (AABB), single (aaBB / AAbb) and double (aabb) *NAC5-1* mutants (n=14 to 24). Average SPAD value at each timepoint was compared against wild type by Wilcoxon test, *=p<0.05; **=p<0.01. (c) Heading date (days after sowing), (d) 25% flag leaf yellowing (DAH), (e) Flag leaf SPAD value of 30 (DAH), (f) 100% peduncle yellowing (DAH), (g) Grain mass (g), (h) Thousand grain weight (g). (c-h) ANOVA with post-hoc Tukey test, formula ∼ Row + Block + Genotype, letters show significance groups at p<0.05. Data from two crosses were combined (n=14 to 24).

To test whether these mutations in *NAC5-A1* and *NAC5-B1* delay senescence, a time-course of flag leaf senescence was measured in TILLING lines at the BC_2_F_3_ generation in the glasshouse. Results show combined data from the two independent crosses, sharing the same mutation in *NAC5-A1* and differing in the mutation in *NAC5-B1*, as both showed similar trends. Wild-type segregants derived from the same crosses were used as controls. On average, *NAC5-1* double mutant lines retained significantly higher flag leaf chlorophyll contents relative to controls from 14-35 days after heading (DAH) (Figure 1b). Similarly, *NAC5-A1* single mutants retained higher chlorophyll contents from 21-42 DAH, and *NAC5-B1* mutants from 21-28 DAH (Figure 1b). Interestingly, all three *NAC5-1* TILLING lines also showed slightly higher chlorophyll contents at heading (Figure 1b). Heading date was earlier in *NAC5-A1* single mutants than controls but did not differ in *NAC5-B1* mutants and double mutants (Figure 1c). The onset of flag leaf senescence was assessed by two methods: a visual score of 25% leaf yellowing and calculation of the point at which the SPAD value passed 30. *NAC5-1* single and double mutant lines trended toward delayed onset of leaf senescence by both metrics, although this was only significant for days to 25% leaf yellowing in *NAC5-A1* single mutant lines (Figure 1 d, e). The time from heading to complete peduncle senescence did not differ between lines (Figure 1f).

To explore whether mutation in *NAC5-1* affects the trade-off between grain mass and protein content, grain traits were also measured in this experiment. Grain mass was significantly higher in *NAC5-A1* single mutants compared to controls but did not differ in *NAC5-B1* mutants or double mutants (Figure 1g). This increase in grain mass may derive from a combination of factors, as neither thousand grain weight, tiller number nor grain number per tiller differed between lines (Figure 1h, Supplementary Figure 4a, b). *NAC5-1* double mutants showed increased grain length (Supplementary Figure 4c). There were no significant differences in grain width, grain area, grain protein content, or plant height (Supplementary Figure 4d-g).

### 3.3 Development of transgenic lines to overexpress *NAC5-A1*

To further assess the effect of *NAC5-1* on senescence timing we expressed *NAC5-A1* from a constitutive rice Actin promoter with a 5’ FLAG tag in hexaploid spring wheat cv. Fielder, selected for its high transformation efficiency (Supplementary Figure 2). Two wheat transformation experiments were carried out with binary construct pMSH30 and transformation efficiencies of 21.8 and 68.4% achieved (percentage of inoculated embryos which regenerate a transformed plant).

To assess the expression of *NAC5-1* in transgenic lines, three sets of qPCR primers were designed to amplify the transgenic construct, the homeolog *NAC5-A1* (including both endogenous and construct-derived transcripts), or all three homoeologs of *NAC5-1* (Supplementary Table 1). Initially, each independently transformed line was assessed by a pooled leaf sample from twelve individuals at the T_1_ generation. As expected, the construct-specific primers amplified in all transgenic lines but did not amplify in non-transformed controls (Supplementary Figure 5a). Based on pooled samples, line 5.4 showed the highest transcript level of *NAC5-A1* among single-copy transformants, while line 8.2 showed the highest transcript level of *NAC5-A1* among multi-copy transformants (Supplementary Figure 5b). Therefore, lines 5.4 and 8.2 were selected. A copy number assay of plants from line 5.4 in the T_1_ generation identified individuals homozygous for the transgenic construct, heterozygous, and wild type segregants (Supplementary Figure 5c).

To account for variability in *NAC5-A1* expression between individual plants, transcript levels were assessed from leaf samples of each plant in the selected lines. Again, the construct was expressed in the majority of plants from transgenic lines but was not amplified in non-transformed controls (Supplementary Figure 5d). Over half of individuals of multi-copy line 8.2 showed a higher transcript level of *NAC5-A1* and of all homoeologs of *NAC5-1* compared to the non-transformed control (Figure 2a, b). On average, homozygous and heterozygous plants of single copy line 5.4 did not show overexpression of *NAC5-A1* or of all homoeologs of *NAC5-1* relative to the non-transformed control (Figure 2a, b). Nevertheless, some individuals within single copy line 5.4 expressed *NAC5-A1* more highly than all non-transformed control individuals (Figure 2a).

**Figure 2.**
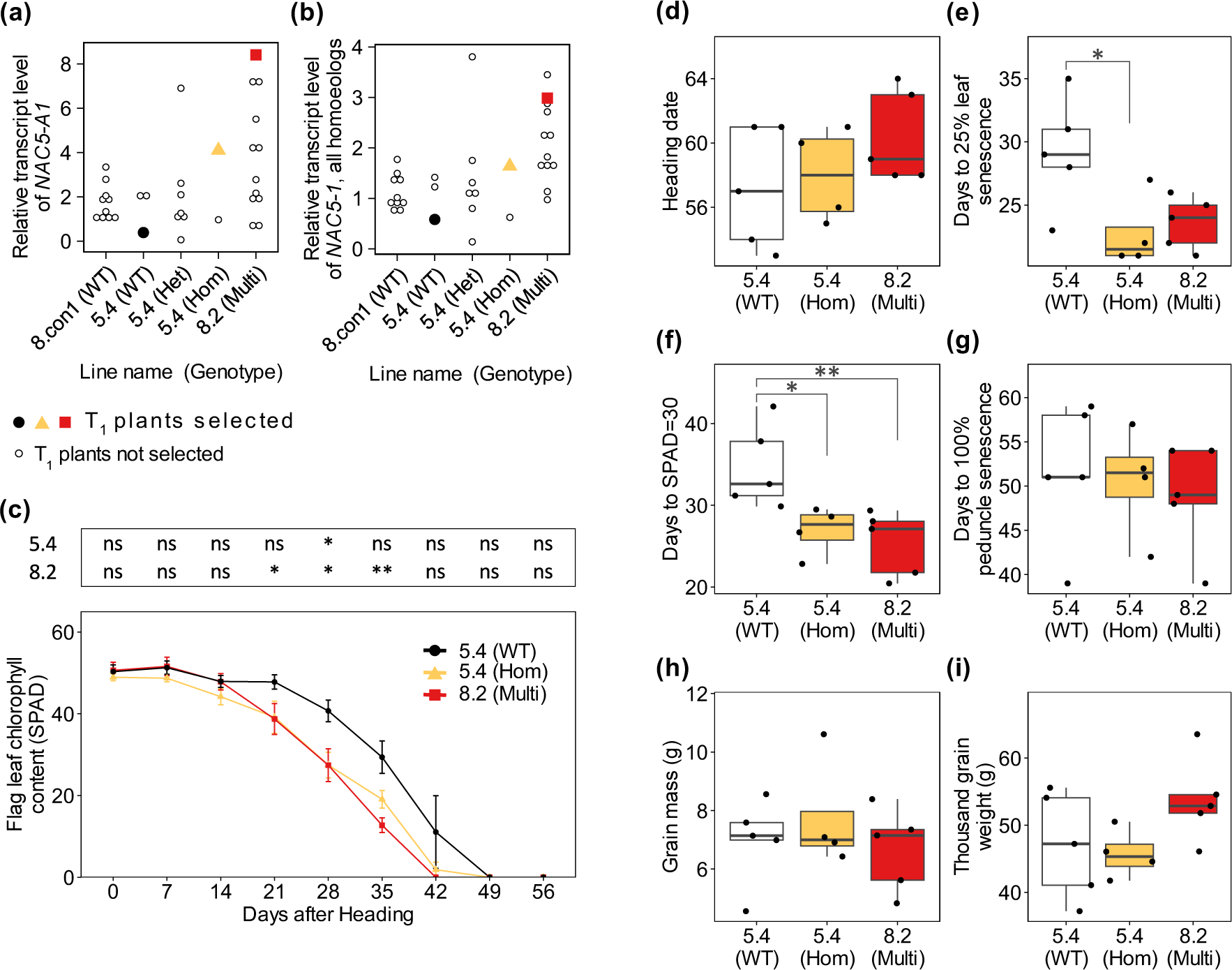
Transgenic lines expressing *NAC5-A1* show earlier onset of flag leaf senescence. (a, b) Relative transcript level of individual 3-week-old T_1_ leaf samples was analysed using the Pfaffl method with primer efficiencies and Actin as reference gene, average of 3 technical replicates normalized against average of single copy plants. Non-transformed control (8.con1), single-copy line (5.4) categorised by construct genotype, and multi-copy line (8.2) are shown. Highlighted points mark T_1_ plants for which T_2_ progeny were used for phenotyping. (a) Relative transcript level of *NAC5-A1*; (b) Relative transcript level of *NAC5-A1*, *NAC5-B1* and *NAC5-D1* combined. (c-i) Phenotypes of T_2_ plants. (c) Flag leaf chlorophyll content (SPAD) in days after heading (DAH) +/− 1 day. (d) Heading date (days after sowing), (e) 25% flag leaf yellowing (DAH); (f) Flag leaf SPAD value of 30 (DAH); (g) 100% peduncle yellowing (DAH); (h) Grain mass (g); (i) Thousand grain weight (g). (c-i) Pairwise comparisons by Wilcoxon test for single-copy transgenic line 5.4 (n=4) and multi-copy transgenic line 8.2 (n=5) against wild-type segregant control from line 5.4 (n=5), ns = not significant; *=p<0.05; **=p<0.01.

If *NAC5-1* regulates senescence, it follows that overexpression of *NAC5-1* would lead to earlier leaf senescence. To test this hypothesis, T_1_ plants were selected to advance to the T_2_ generation for phenotyping according to the following criteria:

1. For overexpression of *NAC5-A1,* from multi-copy line 8.2 the plant with the highest transcript level of *NAC5-A1* was selected (Figure 2a).
2. For slight overexpression of *NAC5-A1,* from single-copy line 5.4, a homozygous plant in T_1_ with a higher transcript level of *NAC5-A1* than non-transformed controls was selected (Figure 2a).
3. As negative control, from single-copy line 5.4 a wild-type segregant control with no amplification of the construct was selected (Figure 2a).

### 3.4 Overexpression of *NAC5-1* leads to slightly earlier leaf senescence

*NAC5-1* overexpression line 5.4 showed reduced flag leaf chlorophyll content at 28 DAH compared to the matched wild-type segregant control, and no difference at other timepoints (Figure 2c). *NAC5-1* overexpression line 8.2 showed a significantly lower average SPAD value from 21-35 DAH than the wild-type control (Figure 2c). *NAC5-1* overexpression lines did not differ from the control in heading date (Figure 2d). Both overexpression lines showed significantly earlier onset of flag leaf senescence scored as days to a SPAD value of 30, and line 5.4 also when scored as day of 25% flag leaf yellowing (Figure 2e, f) The timing of peduncle senescence did not differ between lines (Figure 2g). There were no significant differences in grain mass, thousand grain weight, tiller number, grain length, grain width or grain area, although line 8.2 had fewer grains per tiller than the control (Figure 2h, i; Supplementary Figure 6a-e).

### 3.5 Putative downstream targets of *NAC5-1* include senescence-related genes

To investigate the potential direct downstream target genes of the transcription factor *NAC5-1,* DAP-seq was carried out on *NAC5-A1* and *NAC5-B1* (Gene IDs in Supplementary Table 4)*. NAC5-D1* was omitted because the peptide sequence of the DNA-binding domain is identical between *NAC5-D1* and *NAC5-A1* (Supplementary Figure 3a). All homoeologs of transcription factors *NAM-1* and *NAM-2* (Gene IDs in Supplementary Table 4), previously shown to be associated with senescence timing, were also tested with DAP-seq.

For each transcription factor homoeolog, the number of reads obtained ranged from 55.7M to 94.6M raw read pairs and 14.7M to 24.3M filtered read pairs (Table 2). After filtering, a total of 277 peaks were called across all eight transcription factors, including 39 for *NAC5-A1* and six for *NAC5-B1* (Table 2). *NAM-B1* had both the highest number of input reads and highest number of peaks called (96) indicating that variability in number of peaks called is partly explained by variability in input read depth (Table 2).

**Table 2.**
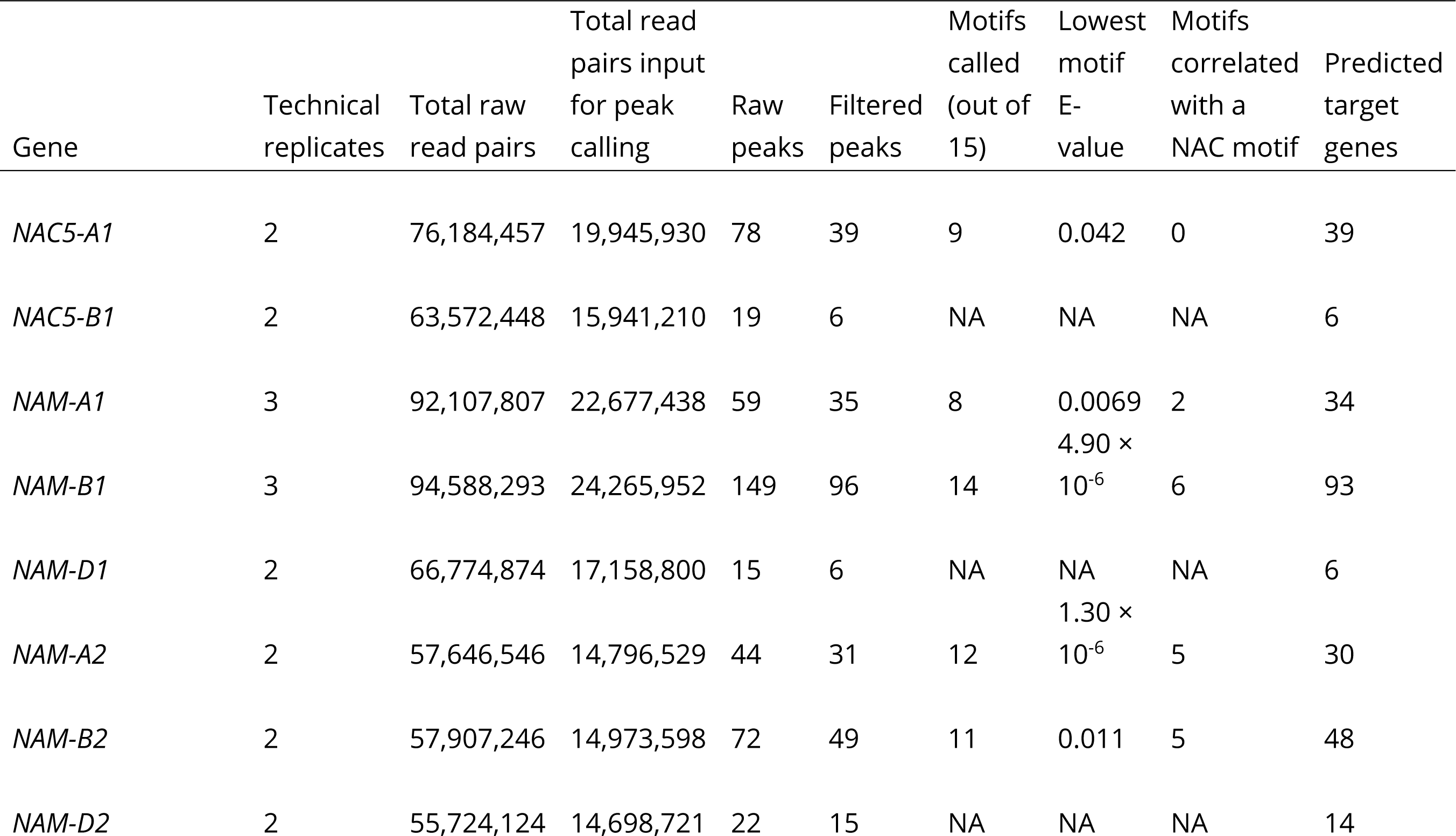
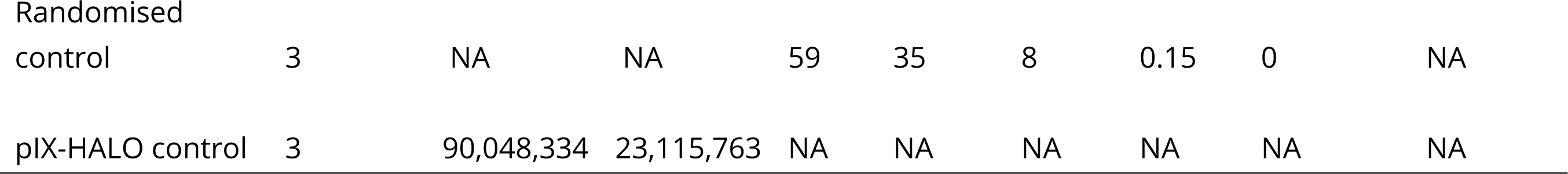
Summary of DAP-seq peaks called for *NAC5-1*, *NAM-1* and *NAM-2*. Peaks were called using two or three pooled technical replicates and filtered for q-value < 0.01 and a greenscreen was applied. Randomised control peak data was generated by randomising location of *NAM-A1* peaks with bedtools shuffle. Motif analysis was run for genes with > 30 peaks.

To assess the quality of the DAP-seq peak data, a motif analysis was carried out on samples with more than 30 peaks. For *NAM-A1, NAM-B1, NAM-A2* and *NAM-D2*, at least two of the *de novo* motifs identified correlated with motifs assigned to NAC family transcription factors in the Jaspar database (Table 2). In addition, the most significant motif for *NAM-B1* and *NAM-A2* showed an E-value less than 1 × 10^-5^ (Table 2). However, for *NAC5-A1* DAP-seq data, and for control data (peaks shuffled to randomised genome locations), none of the motifs identified correlated with NAC transcription factor motifs (Table 2). This indicates that *NAM-A1, NAM-B1, NAM-A2* and *NAM-D2* DAP-seq peak datasets are enriched for transcription factor binding sites, while the *NAC5-A1* dataset may not be.

The closest gene to each DAP-seq peak was identified (Supplementary Dataset 1). The closest genes to peaks in *NAM-A1* and *NAM-B1* shared two genes in common, a higher overlap than would be expected if gene sets were independent (jaccard index 0.016, q<0.01) (Supplementary Dataset 2). Similarly, 11 out of 15 pairwise overlaps between closest genes to peaks in *NAM-A1, NAM-B1, NAM-D1, NAM-A2, NAM-B2* and *NAM-D2* were significant (Supplementary Dataset 2). There was no overlap in closest genes to peaks between *NAC5-A1* and *NAC5-B1,* although an Alpha/beta gliadin gene was shared between *NAC5-A1* and *NAM-B2,* and an uncharacterized gene was shared between *NAC5-B1, NAM-A1* and *NAM-D1* (Supplementary Dataset 3).

These gene sets were also compared to independent datasets for candidate target genes of *NAC5-1*, *NAM-1, NAM-2* and *OsNAC5*. Candidate target genes were obtained from the top 1 million connections of a GENIE3 gene network in wheat (Ramírez-González et al., 2018). Predicted target genes from this gene network showed significant pairwise overlaps between homoeologs within each triad for *NAC5-1, NAM-1* and *NAM-2,* and between *NAM-1* and *NAM-2* (Supplementary Dataset 2). However, while a few genes were common between the gene network and DAP-seq gene sets, none of the pairwise overlaps were significant (Supplementary Dataset 2). Comparison was also made with genes differentially expressed in RNA-seq of *NAC5-A1* overexpression lines compared to wild-type in watered conditions, and identified by DAP-seq in wheat (Mao et al., 2022). Finally, wheat orthologs of candidate target genes of *OsNAC5* in rice based on ChIP-seq and RNA-seq of *OsNAC5* overexpression lines were added (Chung et al., 2018). The genes upregulated in *NAC5-A1* overexpression lines showed a significant pairwise overlap with gene-network predicted targets of *NAC5-A1* and with orthologs of ChIP-seq predicted targets of *OsNAC5* (Supplementary Dataset 2). Overall, 67 genes were associated with *NAC5-1* according to more than one gene set (Figure 3, Supplementary Dataset 3). Of these, 37 were identified as interacting with two or three of the homoeologs of *NAC5-1* within the GENIE3 gene network, while 30 were identified by independent methods in two or more studies (Figure 3, Supplementary Dataset 4). These 67 genes come from a range of different families including cytochrome P450 genes which have been associated with chlorophyll catabolism (Christ et al., 2013) and chloroplast function (Cui et al., 2021), transcription factors in the MYB, bHLH and NAC families which may be part of the senescence regulatory cascade, peptidases which have been associated with protein catabolism in senescence (Roberts et al., 2012) and peroxidases which are known to be upregulated during senescence (Bhattacharjee, 2005).

**Figure 3.**
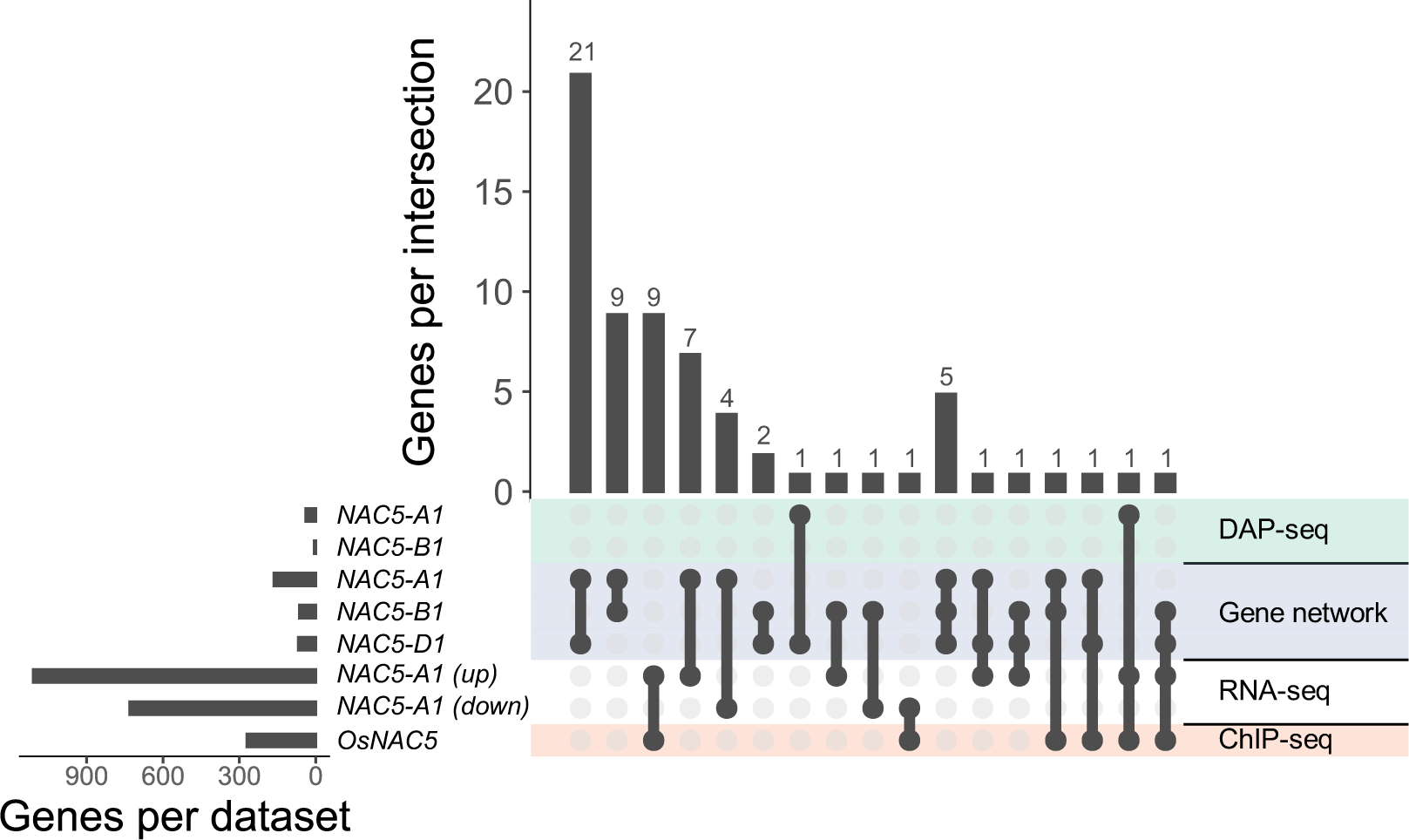
Candidate target genes of *NAC5-1* which overlap in two or more datasets. Left panel shows total number of genes per dataset. Right panel shows the intersection of two or more datasets with black dots, and above, the number of genes in this intersection. DAP-seq: genes closest to peaks in DAP-seq data of *NAC5-A1* and *NAC5-B1*, from this study. Gene network: candidate target genes of *NAC5-A1*, *NAC5-B1* and *NAC5-D1* based on a GENIE3 gene network, from (Ramírez-González et al., 2018). RNA-seq: genes both significantly upregulated (up) or downregulated (down) in RNA-seq of young leaves of *NAC5-A1* overexpression lines in wheat, and identified in a DAP-seq experiment, from (Mao et al., 2022). ChIP-seq: wheat orthologs of rice genes identified from both a ChIP-seq experiment of *OsNAC5*, and differential expression in RNA-seq of roots of *OsNAC5* overexpression lines, from (Chung et al., 2018).

## 4 DISCUSSION

### 4.1 Summary of key findings

We found that mutation of *NAC5-1* was associated with a delay in onset of leaf senescence, while overexpression of *NAC5-1* was associated with earlier leaf senescence, supporting the hypothesis that *NAC5-1* positively regulates the timing of leaf senescence. Thousand grain weight did not differ in *NAC5-1* mutant lines or overexpression lines, and grain protein content did not differ in *NAC5-1* mutant lines, providing no evidence for the hypothesis that by promoting senescence, *NAC5-1* decreases yield and increases protein content. The DAP-seq data generated in this study provided few peaks, and the *NAC5-A1* dataset did not identify a known NAC transcription factor motif, indicating that these data are of limited value. However, comparison with other datasets identified putative downstream target genes of *NAC5-1* which may be involved in senescence.

### 4.2 *NAC5-1* has a conserved role to promote senescence

The retention of higher flag leaf chlorophyll content in *NAC5-1* single and double mutant TILLING lines and more rapid loss of chlorophyll content in *NAC5-1* overexpression lines suggest a role for *NAC5-1* as a positive regulator of the onset of leaf senescence. The chlorophyll retention in *NAC5-1* TILLING lines occurred around 21-35 DAH, coinciding with the upregulation of *NAC5-1* transcripts in wild-type plants at 23-26DAA (Borrill et al., 2019). This is consistent with its ortholog in rice, *OsNAC5,* which was shown to be upregulated in leaf senescence (Sperotto et al., 2009). Grain mass did not differ significantly in *NAC5-1* double mutant lines. However, *NAC5-A1* single mutants showed slightly increased grain mass and *NAC5-1* double mutants showed increased grain length. These results are consistent with rice T-DNA insertion lines with enhanced *OsNAC5* expression, which showed decreased grain length, grain number and kernel weight (Wairich et al., 2023), however overexpression of *OsNAC5* in a separate study resulted in increased grain mass (Jeong et al., 2013). To ascertain the effects of *NAC5-1* on yield components and grain yields larger-scale field experiments will be required. Missense mutations in *NAC5-1* did not affect grain protein content in these experiments, contrasting with rice, where increased *OsNAC5* expression correlated with increased grain protein content (Sharma et al., 2019). Further work is needed to assess grain protein content of *NAC5-1* mutant lines in larger-scale trials in the field, which may result in different findings from our experiments in greenhouse conditions, as has been shown for photosynthetic traits in wheat (Sales et al., 2022). Previously, it was shown that *NAC5-A1* shares a role in promoting drought tolerance with *OsNAC5* (Mao et al., 2022; Takasaki et al., 2010). Our experiments provide evidence that wheat *NAC5-A1* may share further conserved roles with its rice ortholog *OsNAC5,* as it is associated with earlier leaf senescence and decreased grain number and length.

To clarify the extent of the effect of *NAC5-A1* on senescence and effects on grain phenotypes it would be valuable to generate complete null mutants, for example using CRISPR-Cas9. Null mutants may show stronger phenotypic effects than the missense mutants characterised here. Similarly, our transgenic lines showed only a moderate increase in expression and future work using a stronger overexpression promoter could reveal stronger or additional effects. Moreover, our study measuring flag leaf chlorophyll content and peduncle yellowing only captures two aspects of the complex senescence process and additional work will be needed to understand any effects of *NAC5-1* at the biochemical, transcriptomic, cellular and whole plant scales (Borrill et al., 2019; Davies & Gan, 2012; Lim, Kim, & Nam, 2007).

### 4.3 Integrating multiple datasets identified putative downstream targets of *NAC5-1*

This study aimed to identify direct downstream targets of *NAC5-1* using DAP-seq. The 17Gb genome of wheat is significantly larger than the 100Mb genome of *Arabidopsis thaliana,* on which the DAP-seq technique was initially validated (O’Malley et al., 2016). Despite combining technical replicates, it is likely that in this experiment there was insufficient read depth for effective peak calling in wheat. A peak q-value filter of 0.01 was analysed because the more stringent q-value filter of 1 × 10^-10^, used in other studies (Klasfeld, Roulé, & Wagner, 2022), returned very few peaks. Future DAP-seq experiments on polyploid wheat or other species with large genomes should increase the number of reads and maximise mapped read depth by increasing DNA library fragment size (to increase the probability of uniquely mapping a read to a subgenome).

Neverthess, DAP-seq works more efficiently for some transcription factors or transcription factor families than others: previously only 529 of 1,812 (29%) *Arabidopsis* transcription factors and 45 of 189 (24%) wheat transcription factors yielded high quality datasets with *de novo* motifs (O’Malley et al., 2016; Zhang et al., 2022). This variability is not random, as out of transcription factors that did not work on the first attempt, only 10% were recovered on a second attempt (O’Malley et al., 2016). Specifically, DAP-seq of *NAC5-A1* did not identify a known NAC transcription factor motif either in this dataset or in the A-genome donor of wheat (*Triticum urartu* Thumanjan ex Gandilyan), implying that *NAC5-1* may be one of the transcription factors less compatible with DAP-seq (Zhang et al., 2021). As codon usage can impact protein folding, it is possible that some alteration to transcription factor protein folding affected DAP-seq results (Liu, 2020). Further research is needed to optimise DAP-seq to work efficiently with a wider proportion of transcription factors. This would unlock the potential of this technique for research focussed on specific transcription factors, or on developing a detailed regulatory network for a specific trait, such as senescence timing.

Although the DAP-seq for *NAC5-1* was not very informative, adding independent published datasets revealed 67 genes that are putative target genes of *NAC5-1* based on two or more gene sets. Some of these putative target genes have potential associations with senescence. Amongst these, several target genes have associations with nitrogen remobilisation including two serine peptidases that were identified as putative targets of *NAC5-1* from the gene network approach and from downregulation in *NAC5-A1* overexpression lines. One of these serine peptidases was also identified as a target of *NAM-A1* by DAP-seq in this study, suggesting a shared pathway. Serine peptidases are upregulated in senescing wheat flag leaves (Gregersen & Holm, 2007), and protein catabolism by peptidases in the flag leaf is a necessary first step in nitrogen remobilization. These genes may provide a direct mechanism by which *NAC5* and *NAM-1* promote nitrogen remobilisation. *NAC5-1* may also regulate the release of nitrogen from chlorophyll (Christ et al., 2013) through cytochrome P450 genes, of which six were identified as putative targets of two *NAC5-1* homoeologs in the gene network. Remobilised nitrogen may be stored in the form of gliadin in the seeds (Cauvain, 2012), consistent with our identification of an alpha/beta gliadin gene as a putative target of *NAC5-A1* and *NAM-B2* based on DAP-seq in this study and as a target of *NAC5-D1* in the gene network.

Other genes associated with senescence were also identified as putative targets. For example “Senescence regulator S40”, related to *AtS40-3* which is associated with earlier leaf senescence in *Arabidopsis,* was downregulated in *NAC5-A1* overexpression lines and a target of *NAC5-A1* in the gene network (Fischer-Kilbienski et al., 2010). Two closely related heavy metal associated genes were also identified from the gene network, which is interesting given the role of *OsNAC5* in metal ion remobilization (Wairich et al., 2023). Three homoeologs of a wheat peroxidase gene were upregulated in *NAC5-1* overexpression lines, and orthologous to an *OsNAC5* ChIP-seq target. Peroxidases contribute to catabolism of lipids during leaf senescence (Bhattacharjee, 2005). Finally, six of the 67 putative *NAC5-1* targets are transcription factor genes. Investigating these transcription factor gene interactions will aid in building the regulatory network of wheat senescence.

### 4.4 Conclusion

In conclusion, these results indicate that NAC transcription factor *NAC5-1* is a positive regulator of leaf senescence in wheat. Results of one out of two experiments suggest that *NAC5-1* is associated with decreased grain length and grain number. Putative downstream targets of *NAC5-1* feature gene families which have roles in senescence and nitrogen remobilisation, but further research is needed to explore whether *NAC5-1* affects yield or grain protein content in the field. Transcription factors regulating senescence could be targeted to develop wheats with a range of earlier and later flag leaf senescence, to balance yield and protein content.

## AUTHOR CONTRIBUTIONS

CE and PB conceived and designed the research with contributions from SLM, EW and JC. CE, SLM, CS and PB performed the research. CS generated and validated the transgenic wheat lines, PB carried out crossing of TILLING mutants and CE carried out genotyping and phenotypic experiments for TILLING and transgenic lines. CE carried out the DAP-seq with contributions from SLM. CE analysed data including phenotypic and genomic data. CE and PB wrote the paper, and all authors contributed comments to the manuscript. All authors have read and approved the manuscript.

## ACKNOWLEDGEMENTS

The authors thank Dr Ruth Bryant for valuable suggestions during this project and Matthew Hope for assistance with binary construct preparation. We thank the John Innes Centre and University of Birmingham Horticultural Services teams for their support in glasshouse experiments.

