## Supplementary Figures and Tables for "Wheat NAC transcription factor *NAC5-1* is a positive regulator of senescence"

**Supplementary Figures and Tables for Evans *et al*. "Wheat NAC transcription factor *NAC5-1* is a positive regulator of senescence".**

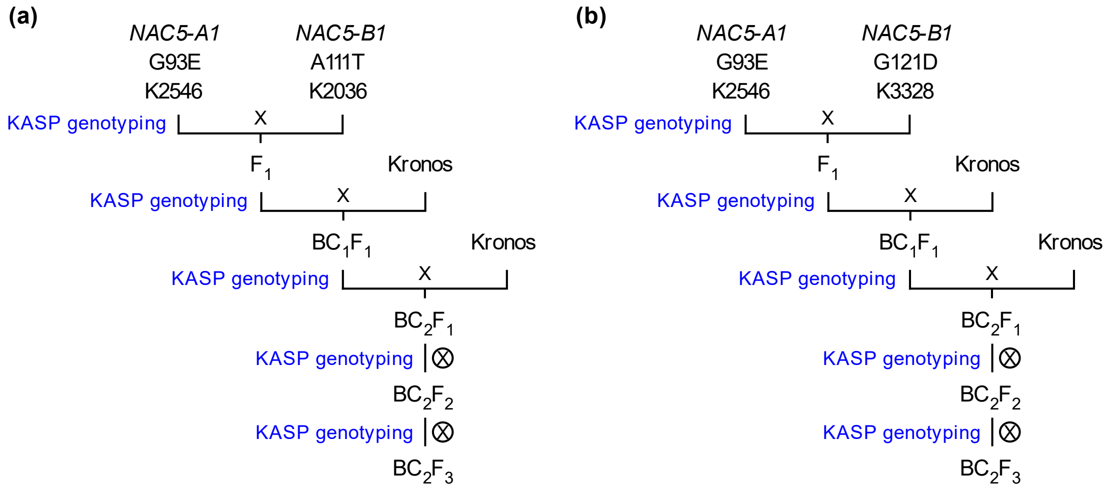


**Supplementary Figure 1.** Crossing scheme for generation of *NAC5-1* TILLING mutant lines. A missense mutation in *NAC5-A1* (K2546) was crossed independently with missense mutations in *NAC5-B1* in (a) K2036 and (b) K3328. Two backcrosses to non-mutagenized Kronos were carried out. Mutations in *NAC5-1* were tracked with KASP genotyping at each generation. $BC_2F_3$ lines were used for senescence phenotyping.

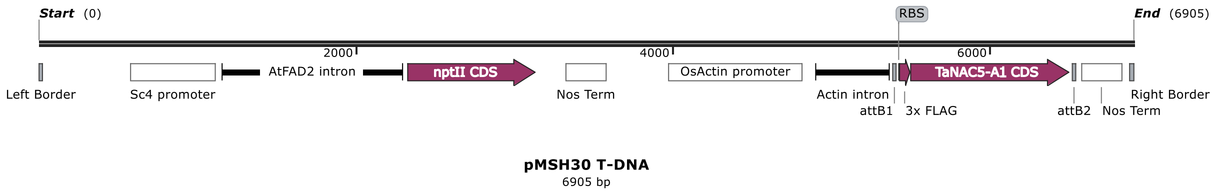


**Supplementary Figure 2.** Schematic of construct for *NAC5-A1* overexpression. The T-DNA region of plasmid pMSH30 includes the *nptII* gene under an Sc4 promoter and *NAC5-A1* with a 5’ 3xFLAG tag under the rice Actin promoter.

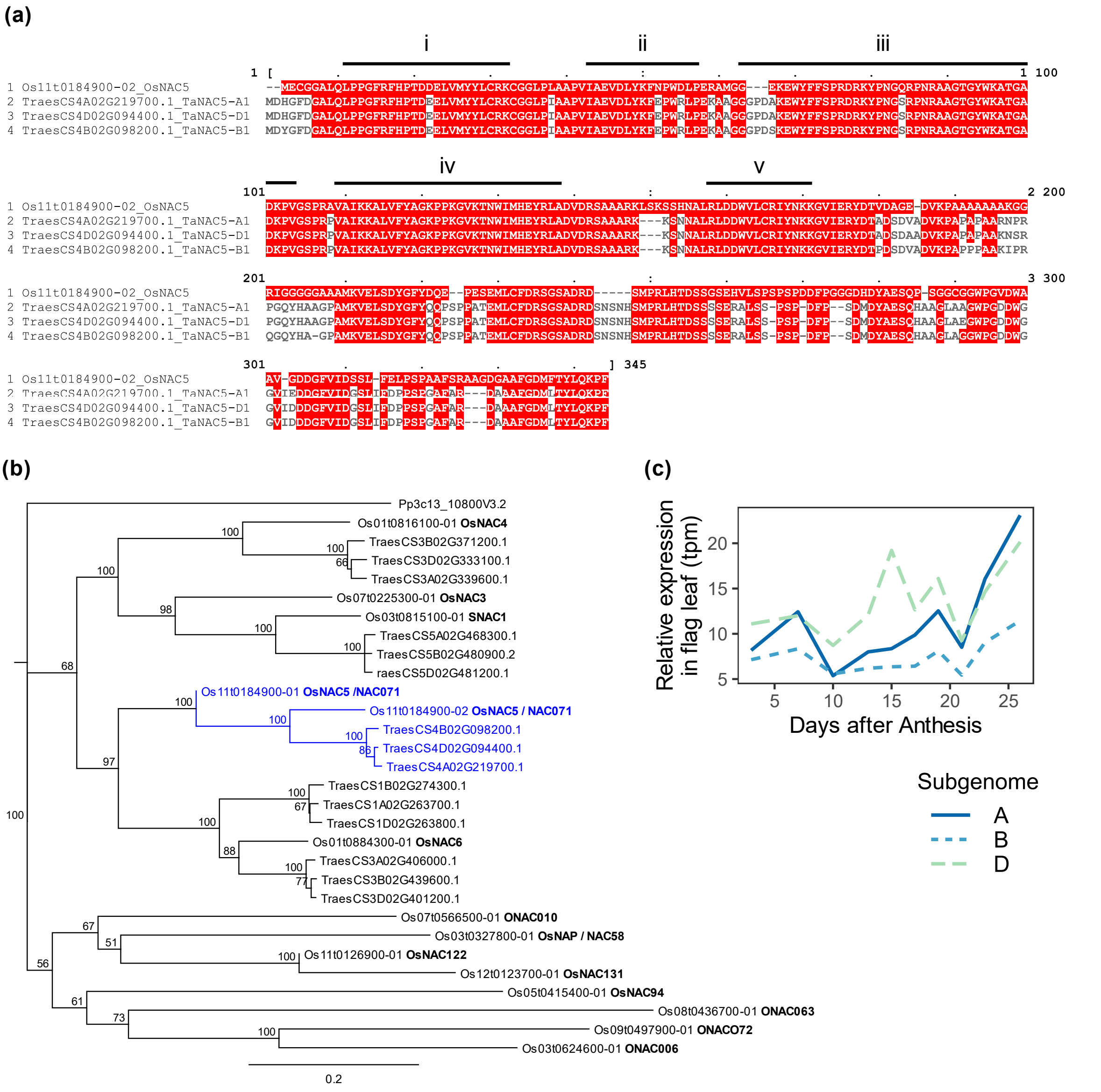


**Supplementary Figure 3.** *TraesCS4A02G219700* and its homoeologs are the orthologs of *OsNAC5* and are expressed in senescence. (a) Alignment of *OsNAC5, NAC5-A1*, *NAC5-B1* and *NAC5-D1* peptide sequences in Clustal Omega. Red marks sites with identity to *OsNAC5* sequence. Conserved NAC subdomains of *OsNAC5* are annotated from (Kikuchi et al., 2000). (b) Peptide sequences of the 15 top BLAST hits of *OsNAC5* in Triticum aestivum and *Oryza sativa* ssp. *japonica* were aligned in Clustal Omega. A rooted tree was built with the Neighbour-Joining method. The node containing *OsNAC5* is highlighted. Numbers show support values from 100 bootstrap replicates. Scale bar shows number of substitutions per site. (c) Expression level of *NAC5-A1, NAC5-B1* and *NAC5-D1* in transcripts per million (tpm) in flag leaves from 3DAA (days after anthesis) to 26DAA. Data from (Borrill et al. 2019).

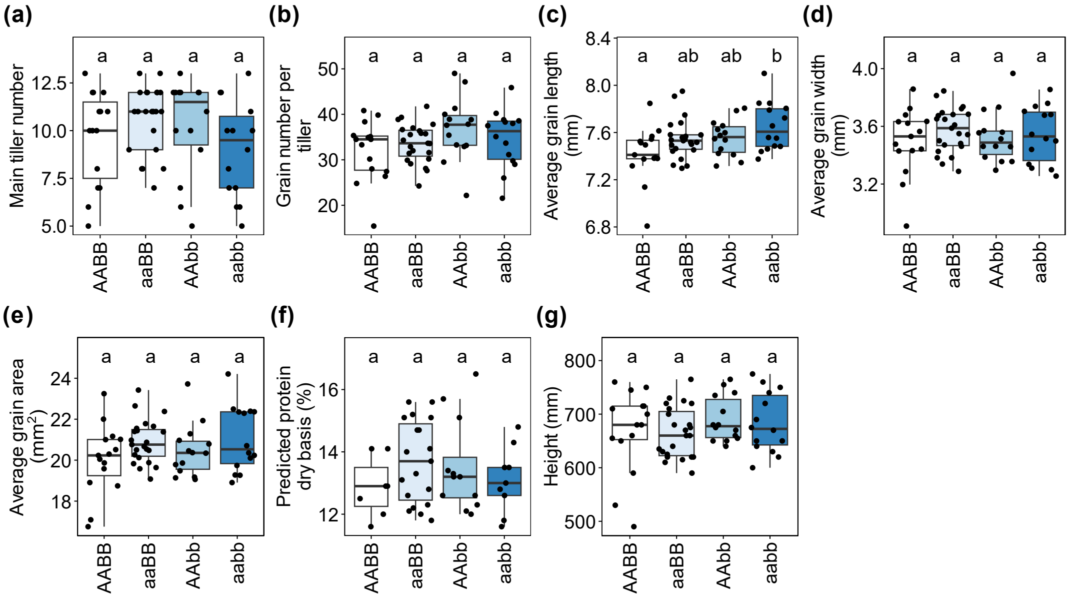


**Supplementary Figure 4.** Additional traits in double mutants of *NAC5-1*. (a) Main tiller number, (b) Grain number per tiller, (c) Average grain length (mm), (d) Average grain width (mm), (e) Average grain area ($mm^2$), (f) Predicted grain protein content by NIR spectrometry, subset of plants with grain mass >15g, dry basis (%), (g) Height of primary tiller (mm). (a-g) ANOVA with post-hoc Tukey test, formula ~ Row + Block + Genotype, letters show significance groups at $p<0.05$. Data from two crosses were combined (n=14-24).

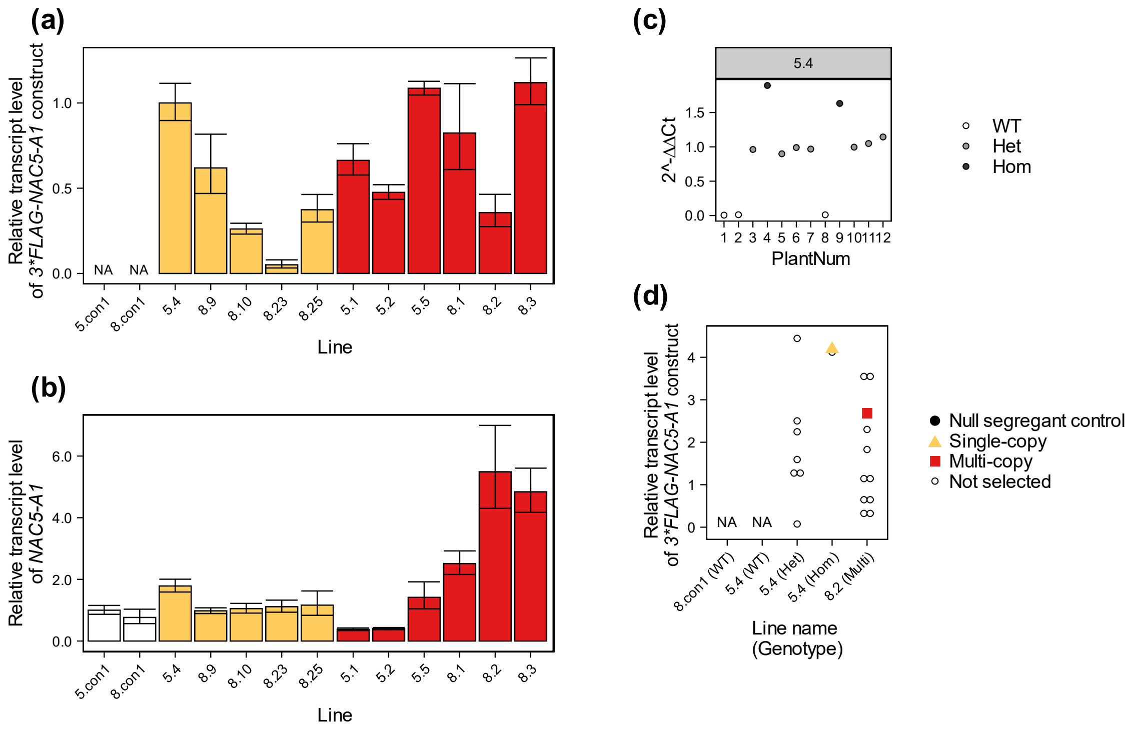


**Supplementary Figure 5.** Selection of *NAC5-A1* transgenic lines. (a, b) Relative transcript level of pooled 3-week-old $T_1$ leaf samples was analysed by qPCR using ΔΔCt method with Actin as reference gene. Bar plot and error bars show mean ±SD (n=3 technical replicates). "NA" marks samples not amplified. (a) Relative transcript level of 3*FLAG::*NAC5-A1* construct normalized against 5.4. (b) Relative transcript level of *NAC5-A1* normalized against 5.con1. (c) Construct genotypes of $T_1$ plants in line 5.4 determined by copy number assay. Relative quantity of genomic DNA of marker gene *nptII* was analysed using Taqman probes and ΔΔCt method, normalised against single copy gene *GAMYB* and average of "Het" cluster. Colours show assigned genotypes. (d) Relative transcript level of individual 3-week-old $T_1$ leaf samples was analysed using the Pfaffl method with primer efficiencies and Actin as reference gene, average of 3 technical replicates normalized against average of single copy lines. Non-transformed control (8.con1), single-copy line (5.4) categorised by construct genotype, and multi-copy line (8.2) are shown. Highlighted points mark $T_1$ plants for which $T_2$ progeny were used for phenotyping.

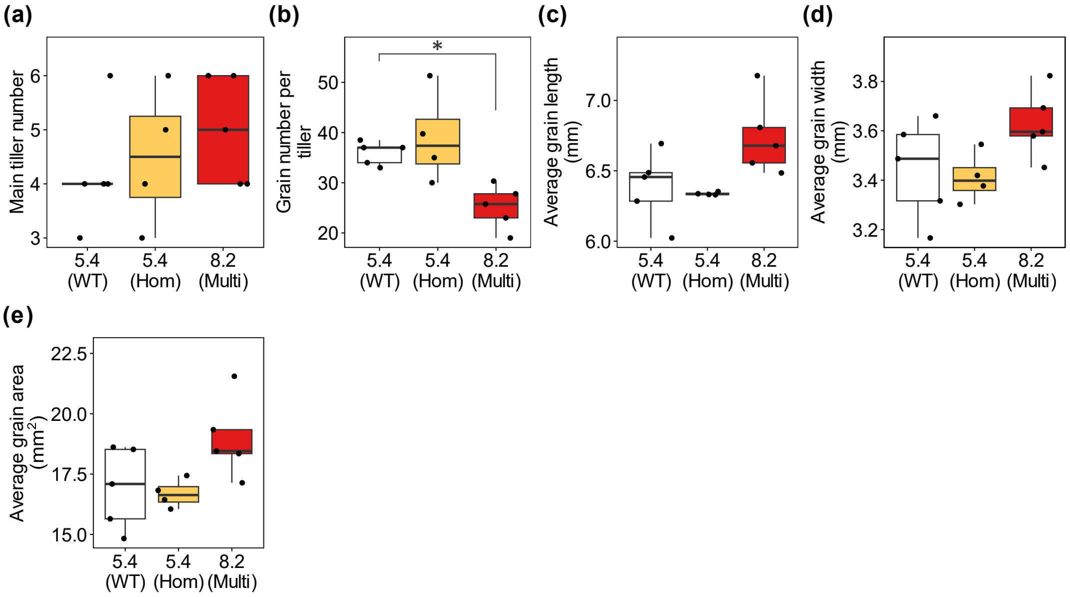


**Supplementary Figure 6.** Further traits in *NAC5-1* $T_2$ transgenic lines. (a) Main tiller number, (b) Grain number per tiller, (c) Average grain length (mm), (d) Average grain width (mm), (e) Average grain area ($mm^2$). (a-e) Pairwise comparisons by Wilcoxon test, ns = not significant; *=p<0.05; **=p<0.01 (n=4-5).

**Supplementary Table 1.** Primers used in this study. The third column corresponds to the Methods section in which these primers were used. Primers sourced from: [1] This study; [2] (Tenea et al., 2011); [3] (Milner et al., 2018); [4] Primers or modification of adapter sequences sourced from NEB.

| Primer name | Sequence | Experiment | Amplifies | Source |
|---|---|---|---|---|
| CENAC5-1-F | GCTCAGGCTGGATGACT | qPCR | *NAC5-A1* | [1] |
| CENAC-7-R | GGCTTGACGTCGGCCA | qPCR | *NAC5-A1* | [1] |
| CENAC5-3-F | TGGATCATGCACGAGTACCG | qPCR | *NAC5-1 (all homoeologs)* | [1] |
| CENAC5-3-R | CACCCAGTCATCCAGCCTG | qPCR | *NAC5-1 (all homoeologs)* | [1] |
| CENAC5-5-F | AAGGACGATGACGATAAGGAC | qPCR | *3*FLAG-NAC5-A1* construct | [1] |
| CENAC5-5-R | GCAAAGGTAGTACATCACCAGC | qPCR | *3*FLAG-NAC5-A1* construct | [1] |
| PB441_Act2_F | CAAATCATGTTTGAGACCTTCAATG | qPCR | *ACT2* | [2] |
| PB441_Act2_R | ACCAGAATCCAACACGATACCTG | qPCR | *ACT2* | [2] |
| PB479_COM | tacttcttctcgccgcgC | KASP | *NAC5-A1* missense in Kronos2546 | [1] |
| PB479_MUT | GAAGGTGACCAAGTTCATGCTcGgtggccttccagtacT | KASP | *NAC5-A1* missense in Kronos2546 | [1] |
| PB479_WT | GAAGGTCGGAGTCAACGGATTcGgtggccttccagtacC | KASP | *NAC5-A1* missense in Kronos2546 | [1] |
| PB480_COM | gggtactggaaggccacG | KASP | *NAC5-B1* missense in Kronos2036 | [1] |
| PB480_MUT | GAAGGTGACCAAGTTCATGCTacgagAgccttcttgatggT | KASP | *NAC5-B1* missense in Kronos2036 | [1] |
| PB480_WT | GAAGGTCGGAGTCAACGGATTacgagAgccttcttgatggC | KASP | *NAC5-B1* missense in Kronos2036 | [1] |
| PB481_COM | gcaaggtggttaccctgagA | KASP | *NAC5-B1* missense in Kronos3328 | [1] |
| PB481_MUT | GAAGGTGACCAAGTTCATGCTtctcgtcttctacgccgA | KASP | *NAC5-B1* missense in Kronos3328 | [1] |
| PB481_WT | GAAGGTCGGAGTCAACGGATTtctcgtcttctacgccgG | KASP | *NAC5-B1* missense in Kronos3328 | [1] |
| GamyB1F | GATCCGAATAGCTGGCTCAAGTAT | Copy number | *GAMYB* | [3] |
| GamyB2R | GGAGACTGCAGGTAGGGATCAAC | Copy number | *GAMYB* | [3] |
| GamyB1P | [Joe]CGTGGCTCCTGCGATGCAGC[TAMRA] | Copy number | *GAMYB* | [3] |
| Npt2B2F | CTCCTGCCGAGAAAGTATCCA | Copy number | *nptII* | [3] |

| | | | | |
|---|---|---|---|---|
| Npt2B4R | GCCGGATCAAGCGTATGC | Copy number | *nptII* | [3] |
| Npt2B2P | [FAM]TGGCTGATGCAATGCGGCG[TAMRA] | Copy number | *nptII* | [3] |
| M13F (-21) | TGTAAAACGACGGCCAGT | DAP-seq | universal vector sequencing | [4] |
| M13R | CAGGAAACAGCTATGAC | DAP-seq | universal vector sequencing | [4] |
| SM_P5_AMP | ACACTCTTTCCCTACACGACGCTCTTCCGATCT | DAP-seq | Illumina Truseq library adapters | [4] |
| SM_P7_AMP | GTGACTGGAGTTCAGACGTGTGCTCTTCCGATCT | DAP-seq | Illumina Truseq library adapters | [4] |
| TaNAC5-A-F | CCACCATGGACTACAAAGACCATGATGGAGAC TATAAGGATCACGACATCGATTACAAGGACGAT GACGATAAGGACCACGGCTTCGACG | Wheat transformation | *NAC5-A1*, adding RBS and 3*FLAG | [1] |
| TaNAC5-A-R2 | TCAGAACGGCTTCTGCAGG | Wheat transformation | *NAC5-A1*, adding RBS and 3*FLAG | [1] |

**Supplementary Table 2.** Summary of copy number analysis of *NAC5-A1* $T_0$ transgenic plants. Copy number analysis was carried out for marker gene *nptII.*

| Copy number | Number of independently transformed plants |
|---|---|
| 1 | 5 |
| 2 | 4 |
| 3 | 1 |
| 4+ | 25 |
| Ambiguous | 8 |
| Non-transformed control | 3 |

**Supplementary Table 3.** Plant growth conditions. Metadata, environmental conditions and experimental design variables (where applicable) are supplied. Watering in all experiments was delivered by automatic flood benching.

| Sowing date | Harvest date | Plant material | Location | Light | Temperature | Pot size | Compost | Other environmental conditions | Experimental layout | Replicates per genotype | Timing of SPAD reading |
|---|---|---|---|---|---|---|---|---|---|---|---|
| 16/11/2020 | 05/03/2021 | *NAC5-A1* overexpression $T_1$ | Glasshouse | 16L:8D Supplementary | 20˚C day/ 18˚C night minimum | 11cm | 3:1 compost: perlite | Standard practice fertiliser and fungicide treatments | NA | NA | NA |
| 23/12/2021 | 20/04/2022 | *NAC5-A1* overexpression $T_2$ | Controlled Environment Chamber | 16L:8D LED 300µMol | 20˚C day/ 15˚C night | 9cm | John Innes Cereal Mix | 70% humidity, 3 fungicide treatments | RCBD | 10 | At Heading, Weekly 6-8DAH to SPAD<10 |
| 01/12/2021 | 19/04/2022 | *NAC5-1* Kronos TILLING | Glasshouse | 16L:8D Supplementary | 20˚C day/ 18˚C night minimum | 1L | John Innes Cereal Mix | Standard practice fertiliser and fungicide treatments | RCBD | 12 | At Heading, Weekly 6-8DAH to SPAD<10 |

**Supplementary Table 4.** Gene IDs used in this study. Gene IDs refer to [1] *Triticum aestivum* assembly IWGSC RefSeq v1.1, [2] NCBI Genbank, or [3] *Oryza sativa* assembly IRGSP-1.0. Gene names coined by [4] This study, [5] (Uauy et al., 2006), [6] (Kikuchi et al., 2000).

| Gene ID | Gene name in this study | Gene ID source | Gene name source |
|---|---|---|---|
| TraesCS4A02G219700 | *NAC5-A1* | [1] | [4] |
| TraesCS4B02G098200 | *NAC5-B1* | [1] | [4] |
| TraesCS4D02G094400 | *NAC5-D1* | [1] | [4] |
| TraesCS6A02G108300 | *NAM-A1* | [1] | [5] |
| DQ869673 | *NAM-B1* | [2] | [5] |
| TraesCS6D02G096300 | *NAM-D1* | [1] | [5] |
| TraesCS2A02G201800 | *NAM-A2* | [1] | [5] |
| TraesCS2B02G228900 | *NAM-B2* | [1] | [5] |
| TraesCS2D02G214100 | *NAM-D2* | [1] | [5] |
| Os11g0184900 | *OsNAC5* | [3] | [6] |
